# Discordance between the predicted vs. the actually recognized CD8+ T cell epitopes of HCMV pp65 antigen and aleatory epitope dominance

**DOI:** 10.1101/2020.11.06.371633

**Authors:** Alexander Lehmann, Ting Zhang, Pedro A Reche, Paul V. Lehmann

**Affiliations:** Cellular Technology Ltd., Shaker Heights, OH, United States; Laboratorio de Inmunomedicina & Inmunoinformatica, Departamento de Immunologia & O2, Facultad de Medicina, Universidad Complutense de Madrid, Madrid, Spain

**Keywords:** Elispot, ImmunoSpot, high throughput, brute force epitope mapping, epitope prediction

## Abstract

CD8+ T cell immune monitoring aims at measuring the numbers and functions of antigen-specific CD8+ T cell populations engaged during immune responses, providing insights into the magnitude and quality of cell-mediated immunity operational in a test subject. The selection of peptides for *ex vivo* CD8+ T cell detection is critical, however, because for each restricting HLA class I molecule present in a human individual there is a multitude of potential epitopes within complex antigens, and HLA diversity between the test subjects predisposes CD8+ T cell responses to individualized epitope recognition profiles. We report here on a brute force CD8+ T cell epitope mapping approach for the human cytomegalovirus (HCMV) pp65 antigen on ten HLA-A*02:01-matched HCMV infected human subjects. In this approach, in each test subject, every possible CD8+ T cell epitope was systematically tested; that is 553 individual peptides that walk the sequence of the HCMV pp65 protein in steps of single amino acids. Highly individualized CD8+ T cell response profiles with aleatory epitope recognition patterns were observed. We compared the actually detected epitope utilization in each individual with epitope prediction ranking for the shared HLA-A*02:01 allele, and for additional HLA class I alleles expressed by each individual. No correlation was found between epitopes’ ranking on the prediction scale and their actual immune dominance. The data suggest that accurate CD8+ T cell immune monitoring might depend on the agnostic reliance on mega peptide pools, or brute force mapping, rather than individualized epitope predictions.

## 1. Introduction

For the past decades, clinical immune diagnostic has relied on the detection of serum antibodies, with interrogation of T cells confined to the scientific discovery. While antibody measurements have become invaluable clinical tests for detecting infections and autoimmune diseases, they have the shortcoming of providing insights into humoral immunity only. There is an increasing number of conditions, however, in which serum antibodies fail to reveal even the immune status of an individual. These include tuberculosis ^1^, Lyme disease ^2^, human immunodeficiency virus (HIV) elite controllers ^3^, up to 30% of human cytomegalovirus (HCMV) infected individuals ^4^,and SARS-CoV-2 infection: in the latter, as with other human coronavirus infections, only a transient antibody response tends to occur, after which specific serum antibodies decline to below detection limit ^5^. In all the aforementioned conditions, however, antigen-specific T cell immunity could be detected (see the corresponding references above), verifying that an in immune response had indeed occurred, suggesting that cell-mediated immunity can occur, and render protection, in the absence of detectable humoral immunity.

Measurement of humoral immunity, therefore, is neither a reliable surrogate for the mere existence of cell mediated immunity, nor does it provide insight into its quality and magnitude. CD4+ and CD8+ T cells and their subpopulations make distinct contributions to host defense however, that, dependent on the pathogen, can be protective or disadvantageous ^6^. This realization has led to an increasing need to progress with T cell immune monitoring, recently precipitated by the SARS-CoV-2 pandemic ^7^.

T cell immune monitoring has a long and successful track record in murine models in which defined experimental conditions, small model antigens, and work with inbred mouse strains expressing few restriction elements (MHC molecules) simplified the task ^8^. The magnitude of scope is entirely different when the outbred human population is to be studied, largely due to the immense diversity in restriction elements (human leukocyte antigens, HLA) and the complexity of the antigenic systems, such as viruses. To comprehensively monitor T cell immunity to SARS-CoV-2, for example, this virus’ entire proteome, 9,871 amino acids long ^9^, would need to be considered. In this report, we confined ourselves to a single protein of HCMV, pp65, which is “only” 561 amino acid long, and to subjects who shared a common HLA-A*02:01 restriction element, but differed in the remaining HLA class I alleles.

Monitoring CD4+ T cell immunity is relatively simple. When the test antigen of interest is added as a protein to peripheral blood mononuclear cells (PBMC), the antigen presenting cells (APC) contained in the PBMC will acquire, process, and present the antigen ^10^. Therefore, natural antigen processing and presentation mechanisms will select which peptide fragments (epitopes) of the antigen will be presented on all HLA class II molecules expressed by an individual. The same spectrum of epitopes that triggered the induction of a CD4+ T cell response *in vivo* will therefore be displayed by the APC in the recall assay *ex vivo*, resulting in the activation of the entire antigen-specific CD4+ T cell repertoire, making sure that no CD4+ T cell epitope will be left behind. For CD4+ T cell immune monitoring, it is therefore not required to know what the potential CD4+ T cell epitopes are in a given donor, nor is there a need to tailor peptides to each test subject’s HLA type – the APC fulfill this function. Unfortunately, this is not the case for *ex vivo* CD8+ T cell detection.

CD8+ T cells evolved to recognize antigens actively bio-synthetized within host cells, as opposed to antigens that APC acquire from their surroundings. Thereby CD8+ T cells can survey ongoing protein synthesis in the cells of the body, permitting them to identify virally-infected or malignant cells, so as to kill them. During protein synthesis, defective byproducts also arise that are degraded by the proteasome into peptide fragments. Such peptides are loaded onto HLA class I molecules, and transported to the cell surface to be displayed to CD8+ T cells ^11^. Protein antigens are not suited to recall *in vivo*-primed CD8+ T cells within PBMC because exogenously added proteins are not efficiently presented to CD8+ T cells in the context of class I molecules. Instead, the CD8+ T cell epitopes need to be added as 8-11 amino acid long peptides that they can bind directly to cell surface expressed HLA class I molecules. From this requirement the need arises to select the “right” peptides for *ex vivo* CD8+ T cell immune monitoring: those very same epitopes that have induced a CD8+ T cell response *in vivo.* Missing the “right” peptides, or only partially covering them, has the consequence that the antigen-specific CD8+ T cell repertoire could go partially or entirely undetected.

Selecting the “right” peptides for CD8+ T cell immune monitoring is an inherently intricate task. HLA class I molecules are encoded by three genetic loci, HLA-A, HLA-B, and HLA-C, for which a multitude of alleles exist in the human population ^12^. Each allelic HLA class I molecule has a unique peptide binding specificity ^13^. As there are barely two humans with an identical HLA-type, there should be barely two humans who present the same array of epitopes. Protecting the species, T cell epitope recognition evolved to be highly individualized ^14^. Peptide selection for comprehensive CD8+ T cell immune monitoring must therefore account for the unique HLA allele composition in each test subject.

A mainstream effort for identifying the “right” peptides for CD8+ T cell monitoring is reliant upon *in silico* epitope predictions. As the peptide binding motifs for most HLA alleles are well-defined, predictions can be made as far as which peptide sequence of an antigen can bind to a given HLA allele, thus constituting a potential T cell epitope. Search engines have been made available to the scientific community to rank peptide sequences for their predicted binding strength to most HLA alleles, thus narrowing in on a finite set of epitopes. A critical assumption for epitope predictions is that peptides that rank high in their respective HLA allele binding score will be those that are being targeted most by CD8+ T cells. The data presented in this study challenge this hypothesis supporting the conclusions reached by Mei *et al*^15^.

Beyond doubt, a peptide needs to be able to bind to an HLA allele to be a candidate for T cell recognition. However, whether a peptide sequence of a protein antigen that has HLA-binding potential indeed becomes an epitope recognized by T cells is defined by many additional factors ^16^. Limitations exist on the level of antigen presentation, including whether that exact peptide is indeed generated through natural antigen processing, and whether it is produced in quantities that can outcompete other peptides, including self-peptides, for binding to the respective HLA molecules. Limitations also exist on the level of the pre-immune T cell repertoire available to engage in antigen recognition. The duration and abundance of epitope presentation will also affect the ensuing CD8+ T cell response, being regulated both by a virus’ replication biology, and the host’s ability to control the virus. Therefore, it can be expected that only a fraction of peptides with HLA class I binding properties will elicit strong CD8+ T cell responses, becoming dominant epitopes. Other presented peptides might induce a weaker, subdominant, barely detectable, cryptic, or no CD8+ T cell responses at all. As all antigen-specific CD8+ T cells can be expected to contribute equally to the host’s defense, irrespective of their fine specificity, comprehensive immune monitoring must not focus on a single or few epitopes, but should instead accommodate all epitopes of an antigen targeted by CD8+ T cells in an individual in order to assess the entire antigen-specific T cell pool.

Next to predictions *in silico*, experimentally verified epitopes have been used as a guide to select peptides for CD8+ T cell immune monitoring. Over the years, T cell lines and clones specific for many viral antigens have been isolated and their epitope specificity compiled in databases ^17, 18^. Selection of such previously verified epitopes for immune monitoring is based on the assumption that epitope recognition, including epitope hierarchy, is constant in subjects who express the corresponding HLA allele. In other words, if an HLA-X-restricted peptide Y has been identified as an immune dominant epitope in an HLA-X positive subject Z, this peptide Y will also be immune dominant in HLA-X positive subjects V and W. Such predictable immune dominance prevails in simple murine models when inbred mice are studied that express minimal restriction element diversity ^19^. However, predictable epitope dominance is lost as soon as restriction element diversity rises through interbreeding these inbred mouse strains. In such F1 mice, aleatory epitope recognition prevails ^20^: T cells in each F1 mouse respond in an unpredictable, dice-like fashion *(alea* means dice in Latin) to epitopes to which the parental strains responded predictably. Aleatory epitope dominance may also apply to humans due to their diverse restriction element makeup ^21^. Therefore, in the present study of HCMV pp65 epitope recognition in HLA-A*02:01-positive individuals, we also compare the peptides that the CD8+ T cells actually target in our cohort with previously verified epitopes.

The third approach for CD8+ T cell immune monitoring is not to select peptides at all, but to systematically test all possible peptides of the antigen. This can be done by using mega peptide pools consisting of hundreds of peptides that cover entire proteins of a virus. By necessity, this approach has become standard recently in the first real world challenge on clinical T cell immune monitoring: trying to study T cell immunity induced by SARS-CoV-2 infection. This crude approach is simple and practical, yet permits comprehensive assessment of the entire expressed antigen-specific T cell repertoire in outbred populations, without requiring customization to HLA types of individuals, but it does not reveal the epitope specificity of the antigen-reactive T cells.

In this study, we applied an agnostic approach in which all possible peptides were tested individually on each subject in a “brute force” high-throughput manner ^22^. The ability to test hundreds, even thousands of peptides individually on a subject is a recent technological advancement. The hurdles that needed to be overcome included limitations in PBMC numbers available from a subject, access to extensive custom peptide libraries, high-throughput-capable T cell assay platforms, and automated data analysis. We have developed and report here large-scale epitope mapping strategies that can be readily adopted even in small academic laboratories operating on tight budgets. Empowered by the ability to experimentally verify CD8+ T cell epitope utilization at the highest possible resolution in the rather well-studied HCMV pp65 T cell immune monitoring model, we set out to compare the epitopes actually recognized with those that are predicted, or assumed to be recognized based on existing data. We draw attention to how incomplete our appreciation of an individual’s expressed epitope space currently is, and suggest that neither epitope predictions, nor reliance on known epitopes suffice, but rather that the agnostic route is best suited for comprehensive CD8+ T cell immune monitoring.

## 2. Materials and Methods

### 2.1 Peripheral Blood Mononuclear Cells (PBMC)

PBMC from healthy adult human donors were from CTL’s ePBMC library (CTL, Shaker Heights, OH, USA). The PBMC had been collected by HemaCare (Van Nuys, CA) under HemaCare’s IRB and sold to CTL concealing the subjects’ identities. The donors’ age, sex, ethnicity, and HLA type are shown in S. Table 1. HLA typing was contracted to the University of Oklahoma Health Science Center (Oklahoma City, OK). The ten subjects for this study were selected according to their HCMV-positive status. The frozen cells were thawed following an optimized protocol ^23^ resulting in viability > 90% for all samples. The PBMC were resuspended in CTL-Test™ Medium (from CTL), developed for low background and high signal performance in ELISPOT assays. The number of PBMC plated into the ImmunoSpot^®^ (ELISPOT) experiments was 3 × 10^5^ PBMC per well.

### 2.2 Peptides and Antigens

553 nonamer peptides, spanning the entire amino acid (a.a.) sequence of the HCMV pp65 protein in steps of single a.a. were purchased from JPT (Berlin, Germany) as a FastTrack CD8 epitope library. These peptides were not further purified following their synthesis, however, individual peptides were analyzed by JPT using LC-MS. The average purity of these peptides was 56%. These peptides were delivered as lyophilized powder with each peptide present in a dedicated well of a 96-well plate, distributed across a total of six 96-well plates. Individual peptides were first dissolved in 50μL DMSO, followed by addition of 200μL of CTL-Test™ Medium so as to generate a “primary peptide stock solution” at 100μg peptide/mL with 20% v/v DMSO. From each of these wells, a “secondary, 10X peptide stock solution” was prepared using a 96-well multichannel pipette, in which peptides were at a concentration of 2μg/mL, with DMSO diluted to 0.4%. On the day of testing, 20μL from each well was transferred “en block,” with a 96-well multi-channel pipette into pre-coated ImmunoSpot^®^ assay plates containing 80μL CTL-Test™ Medium. Finally, 100μL of PBMC (containing 3 × 10^5^ cells) in CTL-Test™ Medium was added resulting in a test peptide concentration of 0.2μg/mL with DMSO present at 0.04% v/v.

UV-inactivated entire HCMV virions (HCMV Grade 2 antigen from CTL) at 10μg/mL was used to recall HCMV-specific CD4 cells. CPI (from CTL) was used as a positive control because, unlike CEF peptides, CPI elicits T cell recall responses in all healthy donors ^24^. CPI is a combination of protein antigens derived from CMV, influenza and parainfluenza viruses, and was used at a final concentration of 6μg/mL in ImmunoSpot^®^ assays.

### 2.3 Human IFN-γ ImmunoSpot^®^ Assays

Single-color enzymatic ImmunoSpot^®^ kits from CTL were used for the detection of antigen-induced IFNγ-producing CD8+ T cells. Peptides or pp65 were plated at the above specified concentrations into capture antibody-precoated assay plates in a volume of 100μL per well. These plates with the antigen were stored in a CO_2_ incubator for less than 1h until the PBMC were thawed and ready for plating. The PBMC were added at 3 × 10^5^ cells/well in 100μL CTL-Test™ Medium followed by a 24h activation culture at 37°C and 9% CO_2_. Thereafter the cells were removed, IFNγ detection antibody was added, and the plate-bound cytokine was visualized by enzyme-catalyzed substrate precipitation. After washing, the plates were air-dried prior to scanning and counting of spot forming units (SFU). ELISPOT plates were analyzed using an ImmunoSpot^®^ S6 Reader, by CTL. For each well, SFU were automatically calculated by the ImmunoSpot^®^ Software using its Autogate™ function ^25^. The data are expressed as SFU per 3 × 10^5^ PBMC, whereby each SFU corresponds to the cytokine footprint of an individual IFNγ-producing T cell ^26^.

### 2.4 Statistical Analysis

As SFU counts follow Gaussian (normal) distribution among replicate wells, the use of parametric statistics is appropriate to identify positive and negative responses, respectively ^27^. The 553 individual peptides of the pp65 nonamer peptide library were tested in single wells. For these peptides, the threshold for a positive response was set at SFU counts exceeding 3 SD of the mean SFU count detected in 18 replicate media control wells, the latter defining the background noise of the test system. This cut off criterion for weak (cryptic) responses renders the likelihood for false positive results at 0.3%. Dominant responses were defined by exceeding 10 SD, and subdominant responses between 5 and 10 SD of the negative control.

### 2.5 HLA-Binding Predictions

We assessed peptide-HLA I presentation by predicting peptide-HLA I binding using HLA I allele-specific profile motif matrices ^28^. We considered that a given peptide binds to a specific HLA I molecule when its binding score ranks within the top 3% percentile of the binding scores computed for 1,000 random 9-mer peptides (average amino acid composition of proteins in the SwissProt database). Peptide binding to experimentally defined HLA-A*02:01 restricted epitopes was predicted using netMHCIpan ^28^ at IEDB analysis resource ^17^, reporting percentile binding score. The lower the percentile binding score the better the binding.

### 2.6. Previously Defined Epitopes

Epitope data for HLA-A2-restricted CD8 cell recognition was obtained from IEDB^17^ with the following search settings: positive response only; host human; MHC I allele, HLA-A*0201, source species HCVM, source antigen: pp65. Only peptides 9 a.a. long were considered.

## 3. Results and Discussion

### 3.1. Experimental Design

We took advantage of the fact that ImmunoSpot^®^ assays require as few as 300,000 PBMC per antigen stimulation condition, and that these assays lend themselves to high-throughput testing and analysis. Utilizing only 173 million PBMC per subject, we therefore could test individually 553 nonamer HCMV pp65 peptides, along with 18 negative control replicate wells to establish the background noise, and the CPI positive control in triplicate.

Nonamer peptides were selected because the peptide binding groove of HLA class I molecules accommodates peptides 8-11 amino acid (aa) in length, with the most common peptide size being 9 aa residues ^10^. In contrast, HLA class molecules present longer peptides ^29^. As such, the usage of nonamer peptides in our assays permitted selective activation of CD8+ T cells. Moreover, because the peptide binding groove of HLA class I molecules is closed on both ends ^10^, it is intolerant for frame shifts. The peptide library was designed therefore to walk the pp65 protein in steps of single a.a. with each nonamer peptide overlapping by 8 a.a. with the previous one (S. Figure 1). Importantly, this approach enabled systematic coverage of every possible CD8+ T cell epitope within the pp65 antigen.

To reduce assay variables, all peptides used in this study were from the same vendor, and were synthetized, stored, dissolved, and tested in the same way. Moreover, all the peptides were tested on each PBMC donor in a single experiment, which rendered the peptides the only assay variable.

Standard IFNγ ImmunoSpot^®^ assays with 24h antigen exposure of PBMC were performed; a time period required for blast transformation and CD8+ T cell activation-driven IFNγ secretion to occur, but too short to permit CD8+ T cell proliferation or differentiation during the cell culture. Thus, we measured at single-cell resolution the frequencies of antigen-specific IFNγ-producing CD8+ T cells as they occurred *in vivo* at isolation of the PBMC. This approach, therefore permitted us to firmly measure within each PBMC sample the number of CD8+ T cells responding to each peptide, and thus to compare the frequencies of peptide-reactive CD8+ T cells to establish epitope hierarchies for each donor. Moreover, adding up all peptide-induced IFNγ SFU permits one to assess the cumulative magnitude of the antigen-specific CD8+ T cell population in each donor, in turn allowing for determination of the relative percentage of antigen-specific CD8+ T cells targeting individual epitopes in each test subject.

Such actual measurements of the epitope-specific CD8+ T cells were then compared to (a) the recognition of published HLA-A*02:01-restricted epitopes, and (b) epitope predictions not only for HLA-A*02:01, an allele that all 10 test subjects shared by design, but also for all other class I molecules expressed by the test subjects.

### 3.2. Highly Variable HCMV pp65 Epitope Recognition Patterns in HLA-A*02:01 Positive Subjects

Eighteen replicate wells containing media alone were included for all 10 individuals in our HCMV positive, HLA-A*02:01 positive cohort in order to firmly establish the background noise of the respective PBMC. The mean and standard deviation (SD) of this negative control is shown for all subjects in Table 1, also specifying the cut off values used for analyzing the peptide-induced SFU counts. The 533 individual pp65 nonamer peptides were also tested on each subjects’ PBMC, and the resulting peptide-induced SFU-counts graded: peptides triggering SFU counts larger than 3 and less or equal to 5 SD over the medium control were considered weakly positive or cryptic (highlighted in beige in Table 1). Of note, with the mean plus 3 SD definition utilized in this study, the chance for a datapoint being false positive was negligible, less than 0.3%. Peptides triggering SFU counts more than 5 and less than or equal 10 SD over the medium background (highlighted in yellow) were labelled subdominant, and peptides eliciting SFU counts exceeding 10 SD over the medium control were called dominant (and are labelled in orange in Table 1). We also introduced a fourth category for peptides that recalled CD8+ T cells in frequencies exceeding 100 SFU/300,000 PBMC, calling them super-dominant epitopes (shown in red in Table 1).

**Table 1.** HCMV ppp65 epitope recognition by CD8+ T cells in HCMV positive, HLA-A*02:01 positive subjects. Ten subjects’ PBMC at 300,000 cells per well were challenged with a library of 553 nonamer peptides that systematically covered all possible CD8+ T cell epitopes of the HCMV ppp65 antigen. An IFN-γ ImmunoSpot assay was performed with the spot forming units (SFU) elicited by each peptide recorded. The mean and SD for 18 negative control media wells, and the cut-off values for the color-coded response categories are specified on the bottom of this Table. Only peptides that induced at least one dominant (orange) or super-dominant (red) recall response in at least one subject are listed. Peptides that have been described as HLA-A*02:01 restricted nonamer epitopes in the literature are highlighted in green.

| Peptides Tested |  | Individual Subjects' CD8+ T Cell Response (SFU per 300,000 PBMC) |  |  |  |  |  |  |  |  |  |
| --- | --- | --- | --- | --- | --- | --- | --- | --- | --- | --- | --- |
| Peptide Name | Sequence | ID 1 | ID 2 | ID 3 | ID 4 | ID 5 | ID 6 | ID 7 | ID 8 | ID 9 | ID 10 |
| pp65:018-026 | ISGHVLKAV | 0 | 2 | 2 | 0 | 7 | 1 | 72 | 11 | 0 | 6 |
| pp65:030-038 | GDPVLPHE | 0 | 2 | 3 | 1 | 2 | 2 | 5 | 9 | 1 | 32 |
| pp65:065-073 | STPCHRGDN | 16 | 0 | 0 | 44 | 0 | 1 | 1 | 5 | 0 | 1 |
| pp65:070-078 | RGDNLQVQ | 2 | 0 | 2 | 2 | 0 | 2 | 84 | 28 | 2 | 5 |
| pp65:095-103 | HNPTRGSIC | 15 | 0 | 20 | 0 | 0 | 1 | 2 | 10 | 1 | 1 |
| pp65:097-105 | PTGRSICPS | 0 | 0 | 41 | 1 | 21 | 0 | 9 | 5 | 0 | 2 |
| pp65:103-121 | CPSQEPMSI | 15 | 0 | 6 | 2 | 1 | 0 | 5 | 11 | 2 | 7 |
| pp65:106-114 | QEPMSIYVY | 14 | 0 | 16 | 5 | 0 | 0 | 2 | 13 | 1 | 3 |
| pp65:107-108 | EPMSIYVYA | 0 | 0 | 17 | 2 | 2 | 0 | 2 | 403 | 1 | 1 |
| pp65:114-121 | YALPLKMLN | 22 | 1 | 7 | 0 | 3 | 2 | 23 | 14 | 0 | 3 |
| pp65:115-123 | ALPLKMLNI | 13 | 0 | 5 | 7 | 2 | 0 | 6 | 22 | 6 | 2 |
| pp65:116-124 | LPLKMLNIP | 71 | 0 | 7 | 14 | 2 | 3 | 5 | 18 | 5 | 2 |
| pp65:119-127 | KMLNIPISIN | 5 | 0 | 6 | 102 | 2 | 1 | 21 | 10 | 0 | 0 |
| pp65:139-148 | HRHLPVADA | 13 | 1 | 1 | 18 | 1 | 1 | 6 | 9 | 5 | 3 |
| pp65:141-149 | HLPVADAVI | 7 | 0 | 1 | 0 | 26 | 0 | 5 | 8 | 0 | 3 |
| pp65:142-150 | LPVADAVIH | 11 | 0 | 2 | 10 | 0 | 0 | 0 | 6 | 10 | 1 |
| pp65:144-152 | VADAVIHAS | 1 | 2 | 5 | 0 | 44 | 1 | 2 | 3 | 3 | 6 |
| pp65:149-157 | IHASGKQMW | 0 | 0 | 525 | 1 | 2 | 0 | 1 | 2 | 2 | 2 |
| pp65:151-158 | ASGKQMWQA | 20 | 0 | 2 | 3 | 0 | 1 | 7 | 6 | 1 | 0 |
| pp65:152-160 | SGKQMWQAR | 23 | 0 | 9 | 7 | 0 | 1 | 10 | 9 | 3 | 1 |
| pp65:155-163 | QMWQARLTV | 1 | 1 | 7 | 1 | 10 | 2 | 33 | 13 | 5 | 0 |
| pp65:175-183 | WKEPDVYIT | 1 | 0 | 2 | 0 | 144 | 0 | 1 | 7 | 0 | 1 |
| pp65:188-196 | FPTKDVALR | 1 | 1 | 1 | 5 | 6 | 1 | 695 | 3 | 13 | 1 |
| pp65:203-211 | ELVCSMENT | 118 | 0 | 0 | 2 | 1 | 1 | 21 | 3 | 7 | 1 |
| pp65:208-216 | MENTRATKM | 1 | 1 | 71 | 7 | 5 | 1 | 0 | 14 | 11 | 3 |
| pp65:221-229 | DQYVKVYLE | 1 | 1 | 7 | 1 | 76 | 0 | 0 | 10 | 6 | 0 |
| pp65:228-236 | LESFCEVDP | 0 | 0 | 2 | 6 | 10 | 1 | 1 | 2 | 0 | 5 |
| pp65:250-258 | VEEDLTMTN | 3 | 0 | 3 | 6 | 13 | 3 | 2 | 2 | 2 | 1 |
| pp65:251-259 | EEDLTMTNR | 0 | 1 | 1 | 2 | 1 | 2 | 107 | 1 | 3 | 2 |
| pp65:262-270 | PFMRPHERN | 0 | 1 | 161 | 5 | 0 | 2 | 6 | 3 | 9 | 1 |
| pp65:267-275 | HERNGFTVL | 0 | 0 | 3 | 0 | 0 | 0 | 6 | 2 | 0 | 46 |
| pp65:270-278 | NGFTVLCPK | 0 | 2 | 10 | 0 | 1 | 0 | 5 | 9 | 7 | 310 |
| pp65:273-281 | TVLCPKNMI | 1 | 0 | 9 | 62 | 0 | 2 | 7 | 2 | 3 | 2 |
| pp65:284-292 | PGKISHIML | 11 | 0 | 0 | 11 | 2 | 7 | 6 | 7 | 10 | 18 |
| pp65:320-328 | LMNGQQJFL | 14 | 2 | 10 | 17 | 1 | 0 | 21 | 2 | 1 | 21 |
| pp65:324-332 | QQIFLEVQA | 343 | 0 | 5 | 3 | 3 | 0 | 6 | 5 | 1 | 8 |
| pp65:325-333 | QIFLEVQAI | 398 | 1 | 6 | 16 | 5 | 1 | 1 | 13 | 0 | 7 |
| pp65:328-336 | LEVQAIRET | 0 | 5 | 810 | 7 | 1 | 1 | 7 | 10 | 5 | 21 |
| pp65:390-398 | EGAAQGGDD | 0 | 0 | 5 | 56 | 0 | 0 | 7 | 13 | 2 | 6 |
| pp65:395-403 | GDDVWVTS | 2 | 0 | 3 | 10 | 0 | 0 | 14 | 98 | 1 | 2 |
| pp65:417-425 | TPRVGGGA | 0 | 0 | 3 | 32 | 0 | 1 | 10 | 2 | 558 | 2 |
| pp65:418-426 | PRVTGGGAM | 1 | 0 | 6 | 6 | 0 | 0 | 6 | 11 | 192 | 0 |
| pp65:430-438 | STSAGRKRR | 3 | 1 | 0 | 89 | 1 | 0 | 34 | 18 | 11 | 7 |
| pp65:431-439 | TSAGRKRRS | 0 | 0 | 8 | 9 | 0 | 1 | 92 | 17 | 3 | 2 |
| pp65:465-473 | EEDTDESD | 0 | 1 | 11 | 54 | 1 | 1 | 5 | 6 | 7 | 7 |
| pp65:482-490 | FTWPPWQAG | 0 | 21 | 1 | 14 | 1 | 1 | 3 | 3 | 10 | 3 |
| pp65:492-500 | LARNLVPMT | 21 | 2 | 5 | 6 | 0 | 0 | 2 | 1 | 5 | 0 |
| pp65:495-503 | NLVPMTVAT | 60 | 303 | 1 | 100 | 97 | 148 | 287 | 674 | 14 | 318 |
| pp65:503-511 | VQGNLKYQ | 2 | 1 | 512 | 1 | 0 | 1 | 5 | 3 | 5 | 0 |
| pp65:511-519 | QEFFWDAND | 2 | 1 | 6 | 9 | 0 | 0 | 8 | 17 | 1 | 95 |
| pp65:512-520 | EFFWDANDI | 0 | 28 | 5 | 10 | 1 | 1 | 8 | 13 | 17 | 61 |
| pp65:513-521 | FFWDANDIY | 1 | 25 | 5 | 6 | 0 | 1 | 2 | 8 | 8 | 100 |
| pp65:514-522 | FWDANDIYR | 0 | 2 | 2 | 1 | 0 | 1 | 10 | 13 | 10 | 44 |
| pp65:521-529 | YRIFAELEG | 2 | 1 | 80 | 5 | 0 | 0 | 28 | 7 | 1 | 5 |
| pp65:524-532 | FAELEGVWQ | 2 | 16 | 6 | 8 | 1 | 7 | 24 | 8 | 11 | 6 |
| pp65:544-552 | QDALPGPCI | 2 | 6 | 5 | 3 | 15 | 5 | 13 | 5 | 2 | 2 |
| Negative Controls and Cut-Off Values For Response Categories | $\bar{x}$ | 1.0 | 0.8 | 4.2 | 3.9 | 3.9 | 1.8 | 6.5 | 8.4 | 5.4 | 3.2 |
| | $\sigma$ | 1.0 | 1.3 | 3.6 | 4.4 | 0.5 | 2.4 | 5.6 | 5.6 | 3.7 | 2.7 |
| | $\bar{x} \pm 3\sigma$ | 3.9 | 4.6 | 14.9 | 17.1 | 5.5 | 8.8 | 23.3 | 25.2 | 16.6 | 11.3 |
| | $\bar{x} \pm 5\sigma$ | 5.8 | 7.2 | 22.1 | 25.9 | 6.5 | 13.5 | 34.4 | 36.3 | 24.0 | 16.8 |
| | $\bar{x} \pm 10\sigma$ | 10.7 | 13.7 | 40.0 | 47.8 | 9.1 | 25.3 | 62.4 | 64.2 | 42.5 | 30.3 |
|  |  | >100 SFU | >100SFU | >100SFU | >100SFU | >100SFU | >100SFU | >100SFU | >100SFU | >100SFU | >100SFU |

Table 1 lists peptides that induced at least one dominant or super-dominant recall response in at least one of the ten test subjects in our cohort. Only for these 56 select peptides of the 553 tested are SFU counts shown for the ten donors. Additionally, a color-coding system was utilized in Table I to delineate whether the peptide recalled a super-, dominant, subdominant, cryptic, or no response in the test subject.

As revealed by the color code at a glance, epitope recognition followed highly individual patterns, that are closer dissected below.

### 3.2. Multiple HCMV pp65 Epitopes are Recognized in Each HLA-A*02:01 Positive Subject

As Table 1 lists only super- and dominant recall responses (>10 SD over background), in S. Table 2 we list 58 additional peptides that induced subdominant recall responses (5-10 SD over the background) in at least one of the ten test subjects in our cohort. At a glance, the color code reveals that these peptides are also recognized in a highly individualized pattern. Peptides that recalled cryptic responses (3-5 SD over background) are not listed individually, but their number is specified for each test subject in Table 2, along with the number of subdominant, and dominant and super-dominant epitopes recognized in each donor. Adding up the number of epitopes in all four categories permits one to establish the cumulative number of CD8+ T cell epitopes recognized in each subject, which varied between 5 and 47 HCMV pp65-derived peptides in this cohort 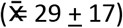. Thus, of the 553 peptides covering the 561 amino acid long pp65 protein, between 1% and 8% 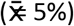 of the peptides constituted a CD8+ T cell epitope in each individual, but for the entire cohort 114 peptides (21% of 553 peptides tested) were needed to recall all dominant (56 peptides) and subdominant (58 peptides) CD8+ T cell epitopes.

**Table 2.** HCMV ppp65 epitope category distribution in HCMV positive, HLA-A*02:01 positive subjects. The number of cryptic, subdominant, dominant and super-dominant epitopes, as defined in the text, are shown for the individual test subjects, along with the sum of epitopes in each category (Total Epitopes Recognized) for each PBMC donor. The absolute number of CD8+ T cells targeting peptides in each category (Cum. SFU) is also shown. From the number of all pp65-specific CD8+ T cells detected in each subject (Cumulative Spec. SFU) the percentage of CD8+ T cells targeting peptides in each of the four response categories has been calculated (% of total SFU). The SFU counts shown are after subtracting the mean + 3 SD specificity cut off value. Because SFU counts for the cryptic category frequently were at the cut-off value, or barely exceeded it, they shown with two decimal places.

|  |  | Test Subjects' CD8+ T Cells Specific for Epitopes |  |  |  |  |  |  |  |  |  |
| --- | --- | --- | --- | --- | --- | --- | --- | --- | --- | --- | --- |
|  |  | ID 1 | ID 2 | ID 3 | ID 4 | ID 5 | ID 6 | ID 7 | ID 8 | ID 9 | ID 10 |
| Cryptic Epitopes |  |  |  |  |  |  |  |  |  |  |  |
|  | Number | 11 | 23 | 32 | 24 | 6 | 2 | 14 | 3 | 8 | 21 |
|  | Cum. SFU | 12.10 | 18.67 | 67.15 | 92.50 | 3.04 | 0.39 | 58.40 | 11.53 | 21.57 | 56.12 |
|  | % of total SFU | 1.02% | 4.41% | 3.02% | 13.54% | 0.73% | 0.24% | 4.03% | 1.04% | 2.89% | 5.09% |
| Subdominant Epitopes |  |  |  |  |  |  |  |  |  |  |  |
|  | Number | 14 | 8 | 5 | 17 | 6 | 2 | 6 | 0 | 1 | 16 |
|  | Cum. SFU | 45 | 35 | 63 | 230 | 14 | 21 | 194 | 0 | 7 | 132 |
|  | % of total SFU | 4% | 8% | 3% | 34% | 3% | 13% | 13% | 0% | 1% | 12% |
| Dominant Epitopes |  |  |  |  |  |  |  |  |  |  |  |
|  | Number | 15 | 4 | 3 | 4 | 9 | 0 | 3 | 1 | 0 | 5 |
|  | Cum. SFU | 281 | 72 | 147 | 193 | 263 | 0 | 178 | 73 | 0 | 221 |
|  | % of total SFU | 24% | 17% | 7% | 28% | 63% | 0% | 12% | 7% | 0% | 20% |
| Super Dominant Epitopes |  |  |  |  |  |  |  |  |  |  |  |
|  | Number | 3 | 1 | 4 | 2 | 1 | 1 | 3 | 2 | 2 | 3 |
|  | Cum. SFU | 847 | 298 | 1948 | 168 | 139 | 139 | 1019 | 1027 | 717 | 694 |
|  | % of total SFU | 71% | 70% | 88% | 25% | 33% | 86% | 70% | 92% | 96% | 63% |
| Total Epitopes Recognized |  | 43 | 36 | 44 | 47 | 22 | 5 | 26 | 6 | 11 | 45 |
| Cumulative Spec. SFU |  | 1185 | 424 | 2226 | 683 | 418 | 161 | 1450 | 1111 | 746 | 1103 |

### 3.3. The Majority of the pp65-Specific CD8+ T Cell Repertoire TargetsSuper-Dominant Epitopes

As T cells recognize processed peptides of antigens there is no reason to assume that a T cell specific for one peptide of the antigen will contribute differently to host defense than T cells recognizing another. Immune monitoring therefore needs to assess all antigen-specific CD8+ T cells irrespective of their fine specificity. In our systematic assessment of CD8+ T cell immunity to pp65, we defined this number as the sum of all SFU counts elicited by the individual epitopes in a subject. This cumulative number of pp65-specific CD8 cells is shown for each subject in Table 2 as “Cum. Spec. SFU”. From this number, one can calculate what percentage of the pp65-specific CD8 + T cells occurs in each of the four response categories. As seen in Table 2, although the number of super-dominant epitopes was low in each subject (between 4 and 1), in eight of ten donors the majority of pp65-specific CD8+ T cells targeted these super-dominant epitopes. The percentage of CD8+ T cells specific for individual dominant and super-dominant epitopes is shown in S. Table 3.

### 3.4. CD8+ T cells Target pp65 Epitopes in an Aleatory Manner

The data in Table 1 and S. Table 2 show that each subject in our cohort displayed a unique CD8+ T cell epitope recognition pattern. This might come as a surprise, as all these subjects were HLA-A*02:01 positive, and one might have expected that among the epitopes recognized there should be at least a shared subset, those restricted by the HLA-A*02:01 allele. To narrow our investigation on such HLA-A*02:01-restricted epitopes, we searched the IEDB database for HLA-A*02:01-restricted nonamer epitopes identifying 31 that have been experimentally verified so far: these are listed in S. Table 4 with the corresponding reference citations. With the exception of the epitope, pp65_495-503_, of all these 31 previously defined HLA-A*02:01-restricted epitopes 5 peptides recalled super-dominant CD8+ T cells responses in only 2 of the 10 test subjects, while 7 additional peptides triggered occasional dominant recall responses. The rest of the 31 previously defined peptides elicited sporadic subdominant (n=4), cryptic (n=6) or no recall responses (n=8) at all. Importantly, donors who did not respond strongly or at all to these previously defined epitopes responded vigorously to other peptides of pp65 (Table 1). These previously defined HLA-A*02:01 -restricted peptides were therefore also targeted in a dice like, aleatory manner in HLA-A*02:01 positive subjects.

Only one previously defined HLA-A*02:01 -restricted epitope, pp65_495-503_, induced a dominant, or super-dominant recall response in 8 of 10 subjects in our cohort (Table 1 and S. Table 4). However, this peptide was not targeted in two donors (ID3# and ID#9) who exhibited responses to other pp65-derived peptides in a super-dominant fashion. Intrigued by this finding, we tested 42 additional (52 in total) HCMV positive, HLA-A*02:01 positive subjects for their recall response to the pp65_495-503_ peptide. As shown in S. Figure 2, the numbers of CD8+ T cells responding to the pp65_495-503_ peptide did not correlate (r^2^ = 0.01) to the numbers of T cells recalled by inactivated HCMV virus; which primarily activates HCMV-specific CD4+ T cells. Even though all these subjects have developed T cell immunity to HCMV, about one fourth of them either did not respond to the pp65_495-503_ peptide, or displayed a low frequency of pp65495-503-specific CD8+ T cells. This finding is consistent with the notion that the CD8+ T cell response to pp65_495-503_ peptide is also aleatory. Interestingly, although the HLA-A*02:01 restriction element was shared by all test subjects in our cohort, and despite the pp65_495-503_ peptide being displayed in vivo via natural processing and presentation, in some individuals this epitope triggered a super-dominant CD8+ T cell response while in other subjects it did not induce a detectable respond at all. Moreover, in yet other donors, the magnitude of the CD8+T cell response induced by this peptide was anywhere between these two extremes. However, the higher prevalence of recognition for this epitope (pp65_495-503_) compared to other previously defined HLA-A*02:01-restricted epitopes might have an unexpected reason: in addition to HLA-A*02:01, pp65_495-503_ received a top binding score for several additional HLA-class I alleles (see next).

### 3.4. HLA Binding Scores are Unreliable Predictors of Actual CD8+ T Cell Epitope Utilization

The participation of the other HLA class I alleles, beyond the shared HLA-A*02:01 restriction element, might explain the highly individual CD8+ T cell response pattern observed in our cohort. Based on extensive knowledge on the peptide binding properties of individual HLA alleles, reference search engines have been established that permit in silico predictions of which peptides fit the binding criteria of a given allele, and moreover, the strength of peptide binding can also be ranked. It has been widely anticipated that such in silico models will suffice to predict epitope utilization. In particular, when there is the need to select one or a few candidate epitopes, e.g. for multimer analysis, it is tempting to pick peptides that have the highest predicted binding score for the HLA allele of interest. In the following we address the validity of such an approach from three angles.

In the first two approaches we focused on predictions for HLA-A*02:01, the most studied HLA allele that is shared by all subjects in our cohort. We introduced into the netMHCIpan ^28^ search engine of the IEDB analysis resource ^17^ the individual sequences of our pp65 nonamer peptide library, resulting in the predicted pp65 ranking shown in Table 3 (in which only the top 30 predicted peptides of 553 are shown). In Approach 1, we compared this *in silico* predicted epitope hierarchy for pp65 with the actual peptide recognition we detected in our cohort. As can be seen in Table 3, pp65_495-503_ ranked as the top binder, and indeed induced a CD8+ T cell response in the majority of our HLA-A*02:01 positive cohort (albeit in an aleatory manner, see above). Most of the other predicted peptides with high HLA-A*02:01 binding scores recalled CD8+T cells in low frequencies, and in an aleatory manner with each of these predicted peptides eliciting SFU in only one or two of the 10 test subjects. Except for the pp65_495-503_ peptide, none of the super-dominant, and few of the dominant responses were recalled by these top 30 HLA-A*02:01 binding peptides (compare with Table 1). One might rightfully argue that this is because those dominant peptides were restricted by, and are binders of alternative class I molecules present in our cohort. We will address this hypothesis below, in Approach 3.

**Table 3.**
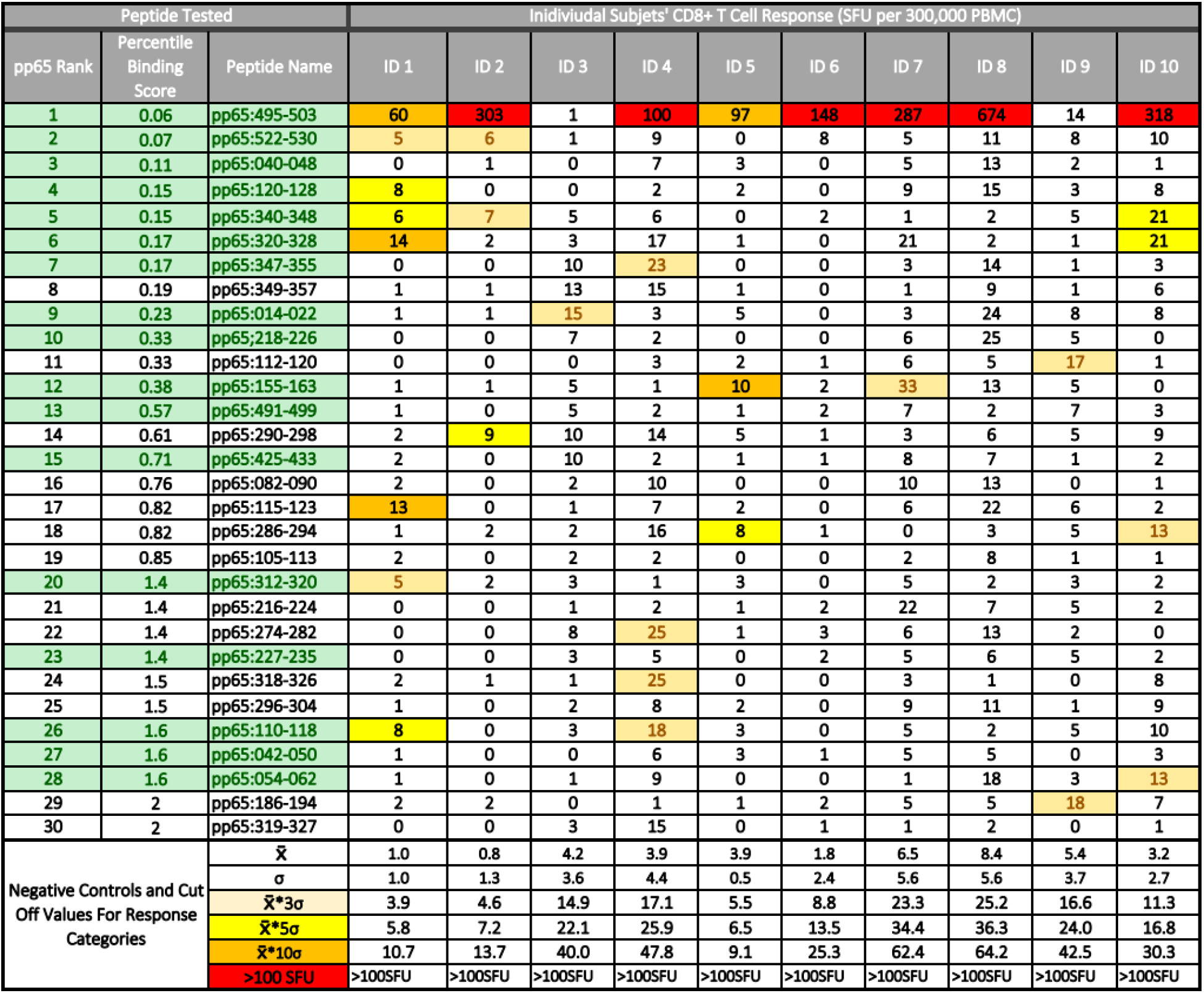
CD8+ T cell recognition of predicted high HLA-A*02:01 -binding peptides. All 553 nonamer pp65 peptides in our library were run on the netMHCIpan search engine of the IEDB Analysis Resource for predicting their binding to the HLA-A*02:01 allele, resulting in “pp65 Rank” shown, with the top binder peptide ranked No. 1. The 30 highest ranking peptides are listed. Additionally, an Percentile Binding Score that compares a peptide’s binding relative to the binding scores computed for 1,000 random nonamer peptides, reporting the percentile binding score, is also listed. A lower percentile binding score denotes better peptide binding to the HLA-A*02:01 allele. Peptides that have been described as HLA-A*02:01 restricted nonamer epitopes in the literature are highlighted in green and the color-coded response categories defined in Table 1 were applied.

In Approach 2, we looked up the predicted HLA-A*02:01 binding scores for those peptides that have been identified experimentally as HLA-A*02:01-restricted pp65 epitopes. In S. Table 4 these peptides have been listed according to their predicted HLA-A*02:01 binding ranking along with CD8+ T cell recall responses they induced in our HLA-A*02:01 positive cohort. With the exception of the pp65_495-503_ peptide, none of these peptides were among the predicted top 20 binders. Seeking for a correlation between the predicted HLA-A*02:01 binding of these peptides, and their actual immune dominance, these data are also represented graphically in S. Figure 3. No significant correlation was seen. The fact, however, that these peptides were targeted by CD8+ T cells in HLA-A*02:01 positive subjects establishes that immune dominant epitopes do not need to rank high in peptide binding score. The score apparently needs to be just high enough to enable stable HLA allele binding.

In Approach 3, we matched binding predictions for all super-dominant and dominant epitopes detected in each of the 10 subjects with all HLA class I alleles expressed in the subject. The results shown for Subject ID 7 in Figure 1 are fully representative for all other subjects in our cohort (see S. Figure 4). As can be seen, the pp65_495-503_ peptide ranked as one of the top binders for several of the subject’s alleles (Figure 1, S. Figure 4), but few of the other actually targeted CD8+ T cell epitopes ranked amongst the top binders for the class I alleles expressed in these respective subjects, and many super-dominant peptides ranked low. A binding score oriented in silico model would not have sufficed to predict the hierarchy of actual epitope recognition.

**Figure 1.**
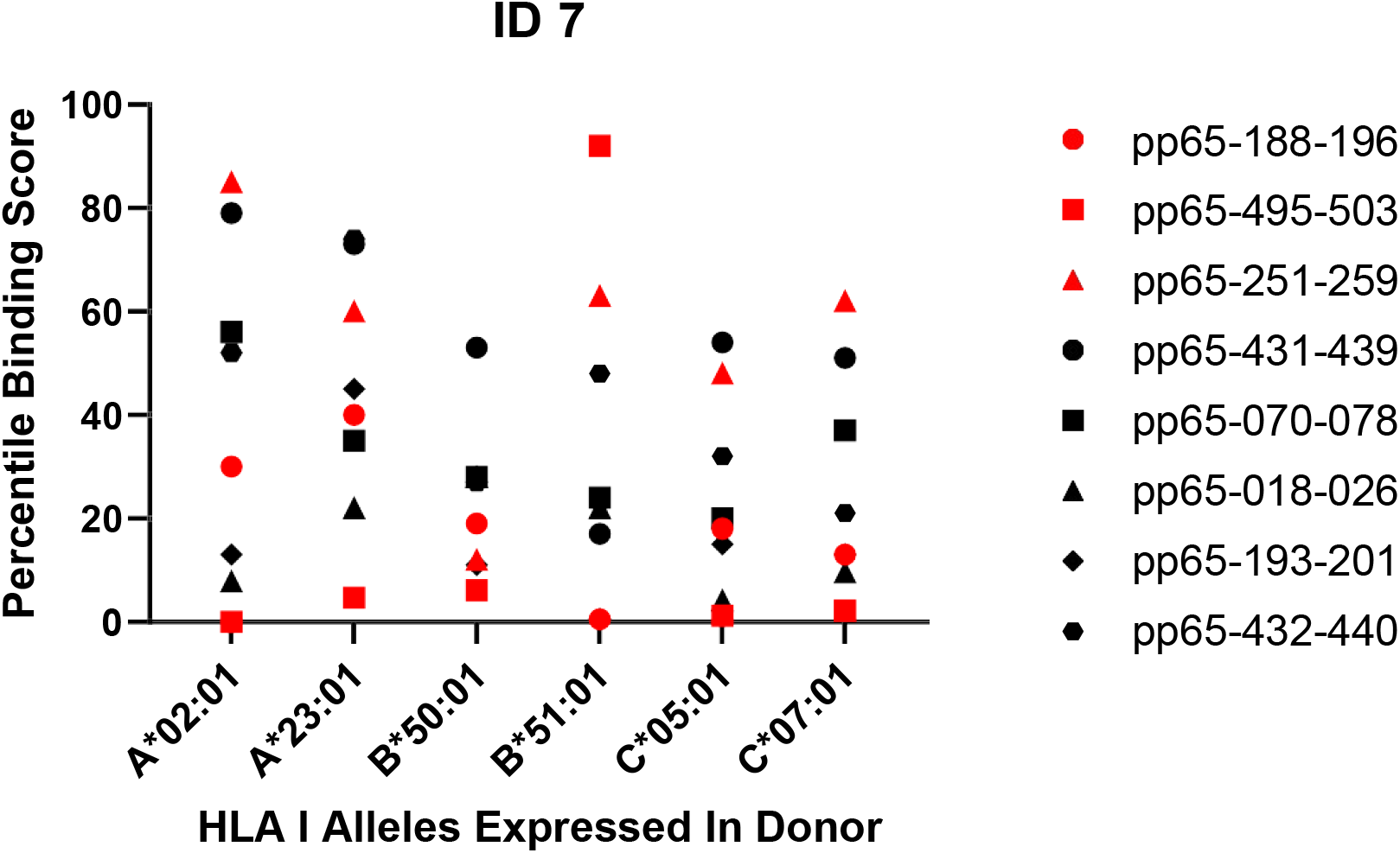
Predicted vs. actual pp65 epitope recognition by CD8+ T cells. Data are shown for subject ID 7 expressing the specified HLA alleles and responding to the listed peptides. Super-dominant responses are shown as red data points, dominant responses in black and weaker responses are not represented. The raw data for the peptide-induced SFU counts are listed in Table 1. The corresponding Percentile Binding Score as established by the netMHCIpan search engine is shown comparing a peptide’s binding relative to the binding scores computed for 1,000 random nonamer peptides. A lower percentile binding score denotes better peptide binding to the specified HLA allele.

All three of our above approaches suggest that, at least in the case of CD8+ T cell immunity induced by HCMV infection against its pp65 antigen, in silico predicted high binding scores for a specific HLA class I allele neither predict whether those peptides will indeed induce a CD8+ T cell response, nor the magnitude of it. This finding raises the question how generalizable it is. Mei *et al*.’s recent report ^15^ suggests that it may be generalizable as they came to the same conclusion studying the prediction performance of databases containing 21,101 experimentally verified epitopes across 19 HLA class I alleles.

## 4 Concluding Remarks

We studied HCMV pp65 antigen recognition by CD8+ T cells in HCMV infected subjects at the highest possible resolution, testing every potential epitope and measuring the numbers of all epitope-specific CD8+ T cells. Our data show that fixed epitope hierarchies do not exist even in an HLA-A*02:01 allele matched cohort. Instead, different super-dominant and dominant epitopes were targeted by the individual test subjects (Table 1). Previously defined epitopes, and peptides predicted to be high HLA-A*02:01 binders also were also targeted in some, but not other individuals, if at all (S. Table 4 and Table 3). If generalizable, the notion of such unpredictable, aleatory epitope recognition patterns in individuals makes it obsolete for CD8+ T cell immune monitoring to rely on testing a select few peptides. Rather, comprehensive CD8+T cell immune monitoring must be all inclusive, accommodating all potential epitopes on all restriction elements of each test subject. Such can be accomplished by brute force epitope mapping, as we did here, or by the use of mega peptide pools.

While in silico epitope ranking may have limited value in predicting immune dominant peptides, it should be helpful for narrowing in on the subset of peptides on an antigen that has sufficient HLA-allele binding affinity to constitute an epitope. As it is impractical to tailor a multitude of variable peptides to each individual’s HLA-type, it might be more realistic for immune monitoring to develop rules for identifying peptides that do not bind to any HLA class I allele, so as to exclude those peptides from testing. Being able to narrow in on peptides should be helpful, as the ultimate goal of immune monitoring is to assess the CD8+ T cell response to the entire proteome of complex antigenic systems, such as all protein antigens of viruses and tumors.

## Acknowledgements

We thank Ruliang Li of Cellular Technology Limited for expert technical assistance, Greg A. Kirchenbaum and Alexey Y. Karulin for valuable discussions, and Diana R. Roen for editorial assistance.

## Data Availability Statement

The datasets generated in this study will be made available by the authors, without undue reservation, to any qualified researcher.

## Author Contributions

Experiments were designed by A.L., PVL and PAR. Experimental data were generated by A.L. with T.Z. contributing.

---

**S. Table 1.** HLA class I allotypes and other characteristics of human subjects tested in this study.

| Code | Ethnicity | Age | Gender | Blood Type | HLA A | HLA B | HLA C |
| --- | --- | --- | --- | --- | --- | --- | --- |
| ID 1 | Hispanic/Latino | 36 | Male | O/Pos | A*02:01 | B*40:08 | C*03:04 |
|  |  |  |  |  | A*68:01 | B*44:03 | C*16:01 |
| ID 2 | Hispanic | 30 | Male | O/Pos | A*02:01 | B*39:05 | C*02:02 |
|  |  |  |  |  | A*24:02 | B*51:01 | C*07:02 |
| ID 3 | Hispanic | 18 | Male | A/Pos | A*02:01 | B*07:02 | C*04:01 |
|  |  |  |  |  | A*02:01 | B*35:17 | C*07:02 |
| ID 4 | Caucasian | 25 | Male | O/Pos | A*02:01 | B*27:05 | C*01:02 |
|  |  |  |  |  | A*23:01 | B*58:01 | C*07:01 |
| ID 5 | Hispanic | 40 | Male | O/Pos | A*02:01 | B*39:05 | C*07:02 |
|  |  |  |  |  | A*68:03 | B*51:01 | C*15:09 |
| ID 6 | Hispanic/Caucasian | 39 | Female | O/Pos | A*02:01 | B*18:01 | C*01:02 |
|  |  |  |  |  | A*25:01 | B*56:01 | C*12:03 |
| ID 7 | Hispanic | 33 | Female | O/Pos | A*02:01 | B*50:01 | C*05:01 |
|  |  |  |  |  | A*23:01 | B*51:01 | C*07:01 |
| ID 8 | Caucasian | 41 | Male | O/Pos | A*02:01 | B*13:02 | C*04:01 |
|  |  |  |  |  | A*23:01 | B*44:03 | C*06:02 |
| ID 9 | Caucasian | 71 | Male | O/Pos | A*02:01 | B*07:24 | C*07:01 |
|  |  |  |  |  | A*03:01 | B*18:01 | C*07:02 |
| ID 10 | Caucasian | 62 | Male | B/Pos | A*02:01 | B*41:02 | C*14:02 |
|  |  |  |  |  | A*66:01 | B*51:01 | C*17:01 |

**S. Figure 1:**
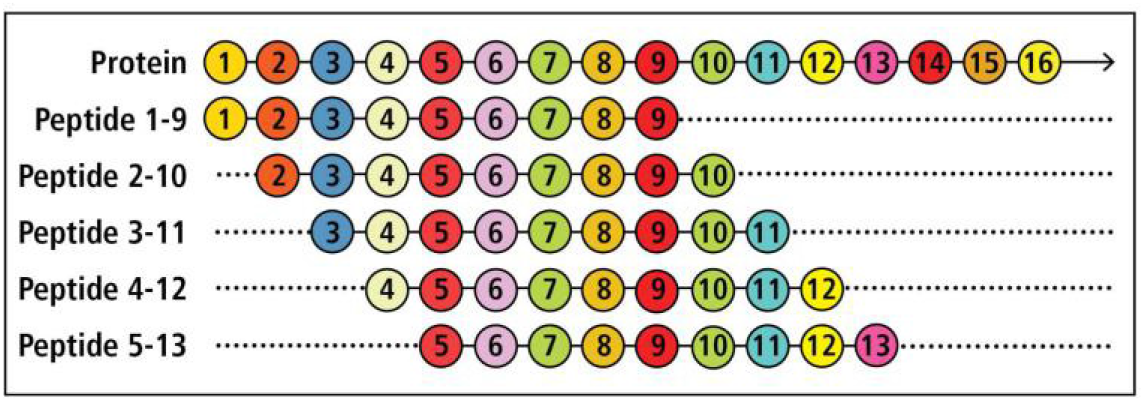
Schematic representation of brute force CD8+ T cell epitope mapping. The amino acid sequence of the protein, illustrated on the top, is covered with nonamer peptides that walk the sequence in steps of single amino acids.

**S. Figure 2:**
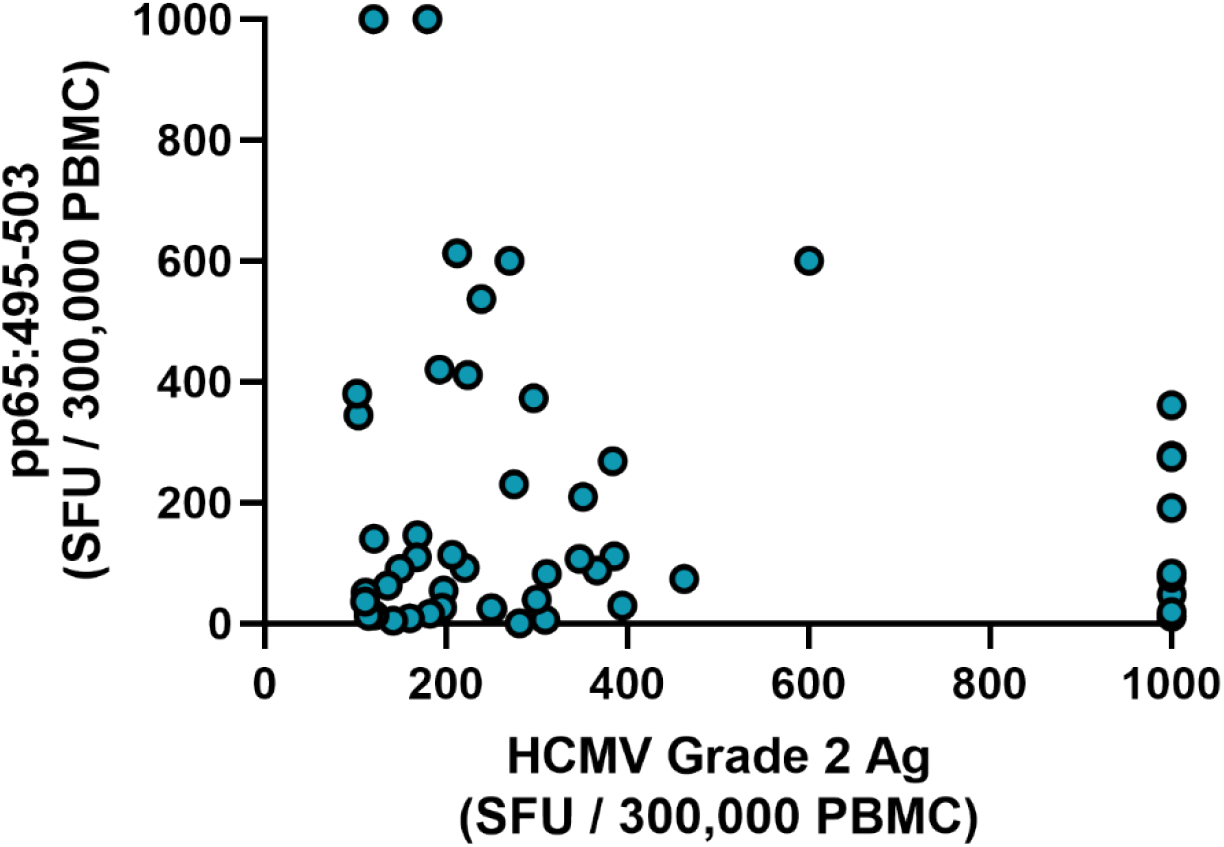
Frequency of HCMV pp65_495-503_ peptide-specific CD8+ T cells vs. HCMV grade 2 antigen-reactive T cells. Fifty-two subjects were selected from the ePBMC database for being HLA-A*02:01 positive and responding to HCMV Grade 2 antigen with more than 100 SFU/300,000 PBMC. Each of these subjects’ PBMC (represented by a dot) were re-tested in an ImmunoSpot assay for the numbers of SFU triggered by HCMV Grade 2 antigen (shown on the X axis), and the numbers of SFU elicited by the pp65_495-503_ peptide (shown on the Y axis).

**S. Table 2:**
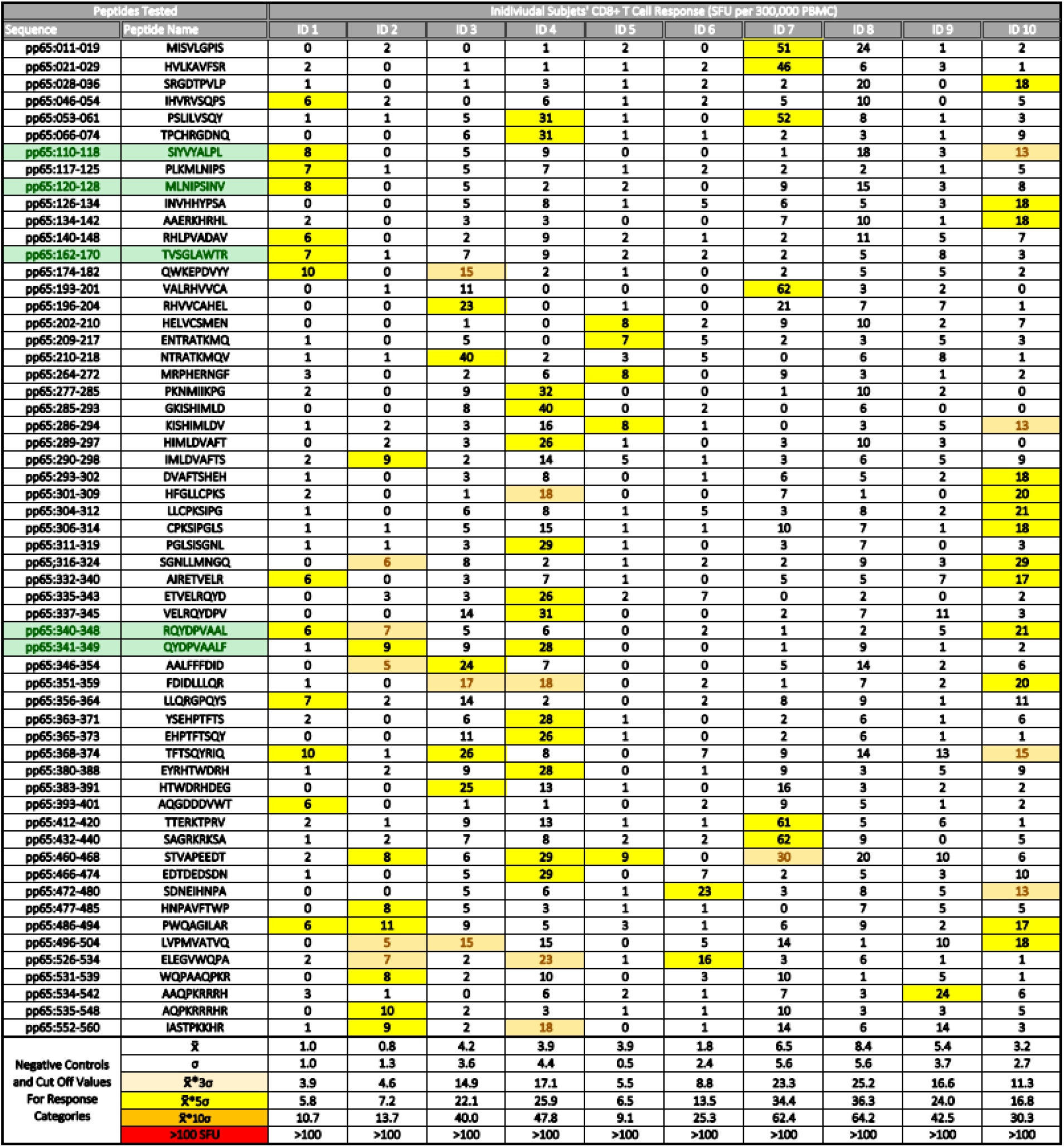
Subdominant pp65 epitopes recognized in our cohort of ten HCMV positive, HLA-A*02:01 positive subjects. Peptides that elicited CD8+ T cell recall responses between 5-10 SD over the media background in at least one of the test subjects are highlighted in yellow. Cryptic recall responses (SD 3-5 over background) triggered by these peptides are highlighted in beige. Peptides that elicited SFU counts exceeding 10 SD over background are not shown in this Table, as they are listed in Table 1. Peptides that only elicited cryptic recall responses (mean plus 3-5 SD) are not shown here, but are summarized in Table 2. Otherwise the legend to Table 1 applies.

**S. Table 3:**
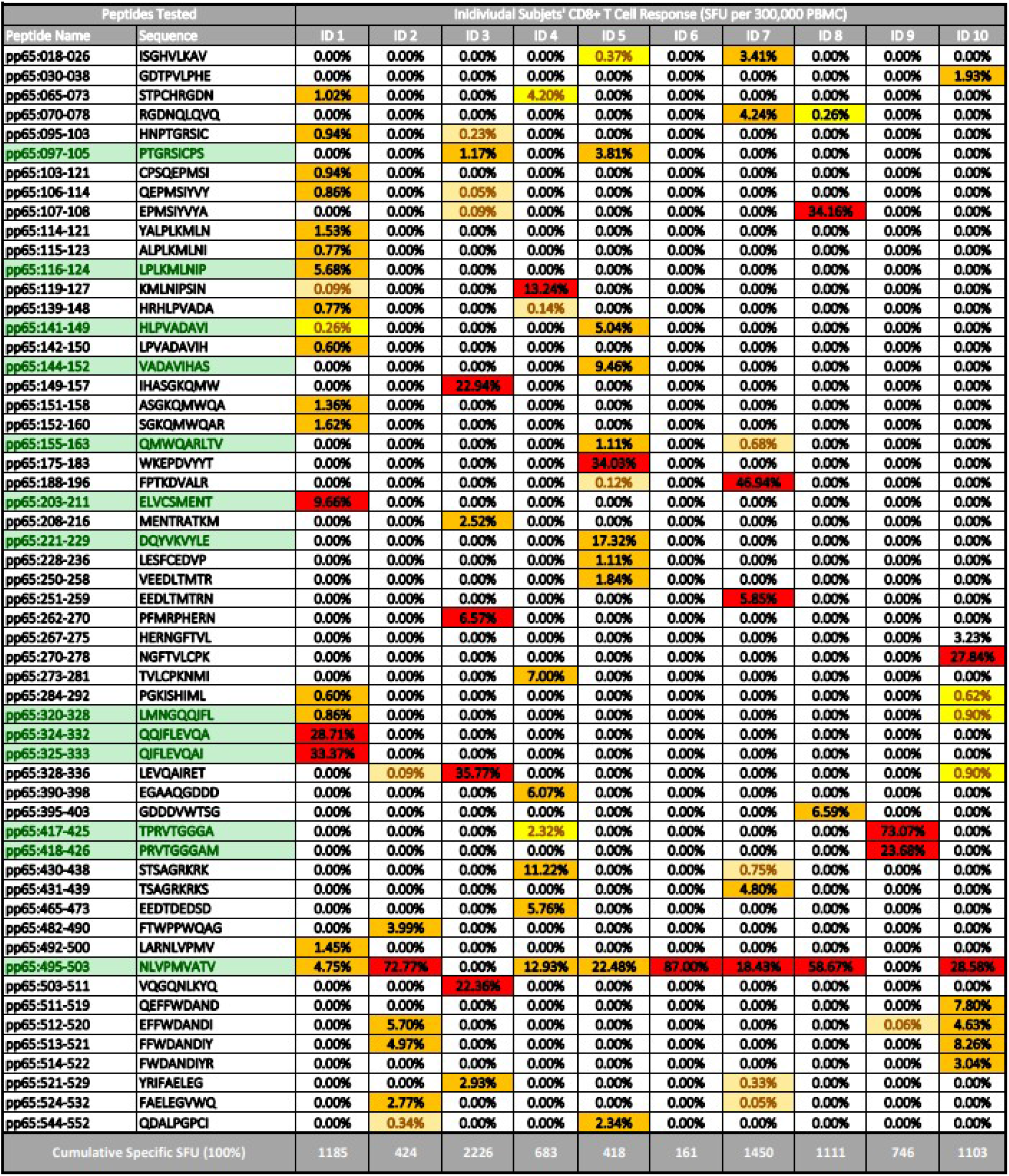
Percentage of pp65-specific CD8+ T cells targeting individual epitopes. The total number of pp65-specific CD8+ T cells was calculated from the sum of SFU triggered by all epitopes in each subject as detailed in Table 2. The percentages of Cumulative Specific SFU counts elicited by individual peptides in each test subjects are shown. For the corresponding absolute SFU counts see Table 1. Otherwise, the legend to Table 1 applies.

**S. Table 4.** Actual recognition of previously identified HLA-A*02:01-restricted nonamer epitopes in our HCMV positive, HLA-A*02:01 positive cohort. The listed peptides have been identified in the IEDB database as HLA-A*02:01-restricted epitopes and the corresponding publications are specified. All 553 nonamer pp65 peptides in our library were run on IEDB’s netMHCIpan search engine for predicting their binding to the HLA-A*02:01 allele, resulting in the “pp65 Rank” shown, with the top binding peptide ranked No. 1. The corresponding Percentile Binding Score is shown comparing each peptide’s binding relative to the binding scores computed for 1,000 random nonamer peptides. A lower percentile binding score denotes better peptide binding to HLA-A*02:01. Otherwise, the legend to Table 1 applies. Following references cited in the table refer to the bibliography 30 22 31 32 33 34 35 36 37 38 39 40 41 42 43 44 45 46 47.

| Ref. | Peptides Tested |  |  |  | Individual Subjects' CD8+ T Cell Response (SFU per 300,000 PBMC) |  |  |  |  |  |  |  |  |  |
| --- | --- | --- | --- | --- | --- | --- | --- | --- | --- | --- | --- | --- | --- | --- |
|  | Peptide Name | Epitope Sequence | Percentile Binding Score | pp65 Rank | ID 1 | ID 2 | ID 3 | ID 4 | ID 5 | ID 6 | ID 7 | ID 8 | ID 9 | ID 10 |
| 42,43,44,45 | pp65:495-503 | NLVPMTATV | 0.06 | 1 | 60 | 303 | 1 | 100 | 97 | 148 | 287 | 674 | 14 | 318 |
| 33 | pp65:340-348 | RQYDPVAAL | 0.06 | 2 | 6 | 7 | 5 | 6 | 0 | 2 | 1 | 2 | 5 | 21 |
| 30,37 | pp65:040-048 | RLQTGIHV | 0.09 | 3 | 0 | 1 | 2 | 7 | 3 | 0 | 5 | 13 | 2 | 1 |
| 30,36,37,38,39 | pp65:522-530 | RIFAELEGV | 0.11 | 5 | 5 | 6 | 0 | 9 | 0 | 8 | 5 | 11 | 8 | 10 |
| 31,37 | pp65:320-328 | LMNGQQIFL | 0.15 | 4 | 14 | 2 | 10 | 17 | 1 | 0 | 21 | 2 | 1 | 21 |
| 30 | pp65:218-226 | VIGDQYVKV | 0.23 | 7 | 0 | 0 | 10 | 3 | 2 | 1 | 6 | 5 | 17 | 1 |
| 31,32 | pp65:155-163 | QMWQARLTV | 0.24 | 6 | 1 | 1 | 7 | 1 | 10 | 2 | 33 | 13 | 5 | 0 |
| 40,41 | pp65:014-022 | VLGPISGHV | 0.24 | 9 | 1 | 1 | 10 | 3 | 5 | 0 | 3 | 24 | 8 | 8 |
| 30,34,37,40,41,46,47 | pp65:120-128 | MLNIPSINV | 0.25 | 11 | 8 | 0 | 5 | 2 | 2 | 0 | 9 | 15 | 3 | 8 |
| 33 | pp65:347-355 | ALFFFDIDL | 0.54 | 12 | 0 | 0 | 8 | 23 | 0 | 0 | 3 | 14 | 1 | 3 |
| 37 | pp65:491-499 | ILARNLVPM | 0.74 | 13 | 1 | 0 | 5 | 2 | 1 | 2 | 7 | 2 | 7 | 3 |
| 30 | pp65:425-433 | AMAGASTSA | 0.85 | 15 | 2 | 0 | 5 | 2 | 1 | 1 | 8 | 7 | 1 | 2 |
| 34 | pp65:042-050 | LQTGIHVRV | 0.98 | 21 | 1 | 0 | 6 | 18 | 3 | 0 | 5 | 2 | 5 | 10 |
| 30 | pp65:054-062 | SILVSQYT | 1.5 | 23 | 1 | 0 | 1 | 6 | 3 | 1 | 5 | 5 | 0 | 3 |
| 22 | pp65:325-333 | QIFLEVQAI | 1.6 | 27 | 398 | 1 | 6 | 16 | 5 | 1 | 1 | 13 | 0 | 7 |
| 30 | pp65:312-320 | GLSISGNLL | 1.9 | 26 | 5 | 0 | 9 | 5 | 0 | 2 | 5 | 6 | 5 | 2 |
| 30 | pp65:110-118 | SIYVYALPL | 2.2 | 28 | 8 | 0 | 5 | 9 | 0 | 0 | 1 | 18 | 3 | 13 |
| 30 | pp65:227-235 | YLESFCEVD | 2.6 | 32 | 0 | 0 | 1 | 2 | 1 | 2 | 22 | 7 | 5 | 2 |
| 34,35 | pp65:341-349 | QYDPVAALF | 3.3 | 36 | 1 | 9 | 9 | 28 | 0 | 0 | 1 | 9 | 1 | 2 |
| 22 | pp65:324-332 | QQIFLEVQA | 4 | 43 | 343 | 0 | 5 | 3 | 3 | 0 | 6 | 5 | 1 | 8 |
| 30 | pp65:519-527 | DIYRIFAEI | 4.3 | 49 | 1 | 0 | 7 | 0 | 2 | 1 | 14 | 3 | 1 | 1 |
| 22 | pp65:141-149 | HLPVADAVI | 5.1 | 54 | 7 | 0 | 1 | 0 | 26 | 0 | 5 | 8 | 0 | 3 |
| 22 | pp65:144-152 | VADAVIHAS | 11 | 80 | 1 | 2 | 5 | 0 | 44 | 1 | 2 | 3 | 3 | 6 |
| 30 | pp65:509-517 | KYQEFFWDA | 12 | 92 | 0 | 2 | 2 | 2 | 0 | 3 | 11 | 2 | 2 | 7 |
| 33 | pp65:345-353 | VAALFFFDI | 16 | 97 | 0 | 6 | 2 | 23 | 0 | 0 | 8 | 3 | 2 | 5 |
| 22 | pp65:203-211 | ELVCSMENT | 23 | 163 | 118 | 0 | 0 | 2 | 1 | 1 | 21 | 3 | 7 | 1 |
| 22 | pp65:221-229 | DQYVKVYLE | 25 | 229 | 1 | 1 | 7 | 1 | 76 | 0 | 0 | 10 | 6 | 0 |
| 22 | pp65:116-124 | LPLKMLNIP | 51 | 360 | 71 | 0 | 7 | 14 | 2 | 3 | 5 | 18 | 5 | 2 |
| 22 | pp65:417-425 | TPRVTGGGA | 63 | 378 | 0 | 0 | 3 | 32 | 0 | 1 | 10 | 2 | 558 | 2 |
| 22 | pp65:418-426 | PRVTGGGAM | 73 | 394 | 1 | 0 | 6 | 6 | 0 | 0 | 6 | 11 | 192 | 0 |
| 22 | pp65:097-105 | PTGRSICPS | 78 | 510 | 0 | 0 | 41 | 1 | 21 | 0 | 9 | 5 | 0 | 2 |

**S. Figure 3.**
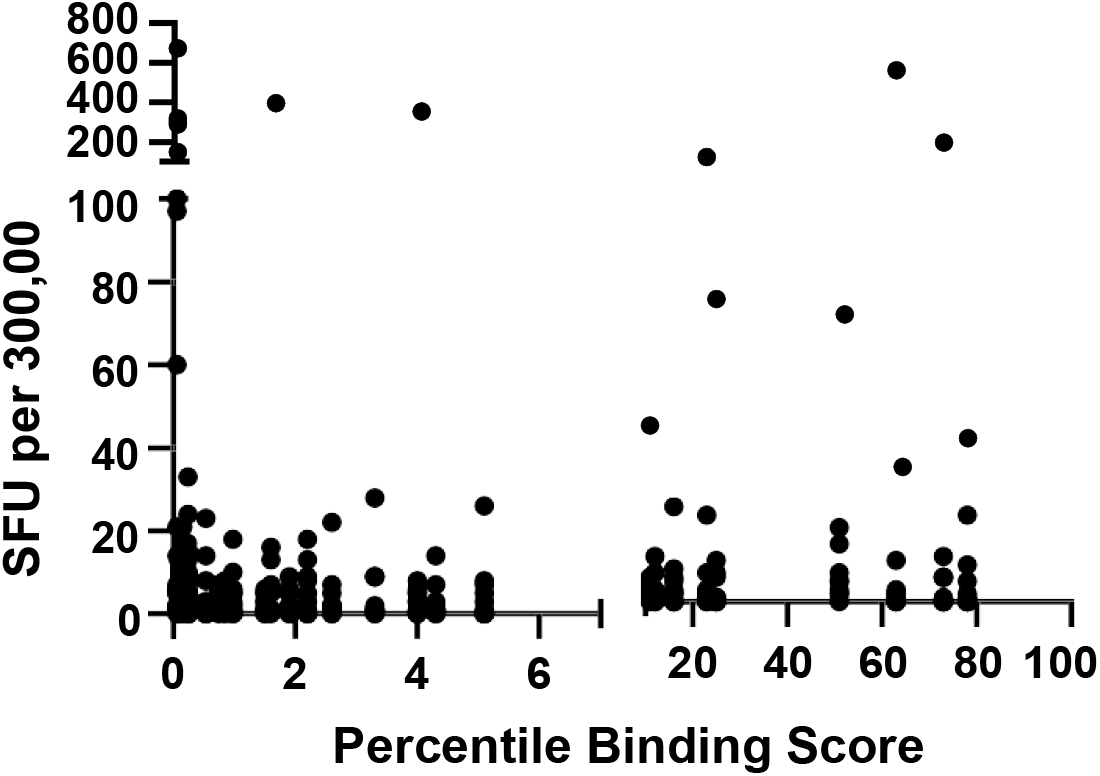
HLA-A*02:01 binding ranking of previously defined HLA-A*02:01 restricted nonamer pp65 peptides vs. the SFU counts they induced in our cohort of HCMV positive, HLA-A*02:01 positive subjects. The numeric SFU data shown in Supplementary Table 4 are plotted relative to their Percentile Binding Score as established run on the netMHCIpan search engine for predicting their binding to the HLA-A^*^02:01 allele.

**S. Figure 4.**
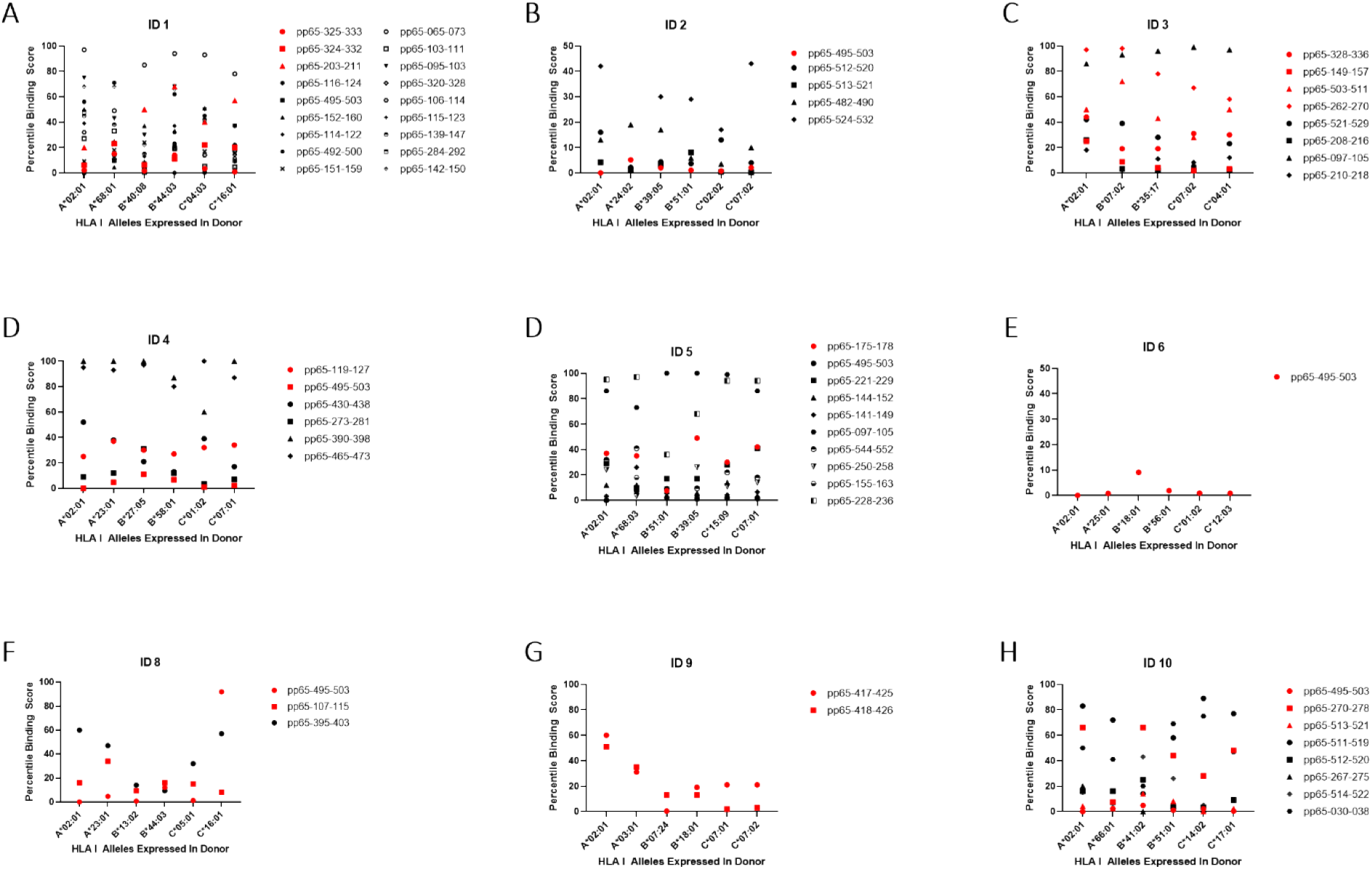
Predicted vs. actual pp65 epitope recognition by CD8+ T cells. Data are shown for the subjects specified in each panel. For each subject his/her HLA class I alleles are specified (in the case of homozygosity the allele is listed once). Peptides that induced super-dominant responses in that individual are shown as red data points, dominant responses in black and weaker responses are not represented. The raw data for the peptide-induced SFU counts are listed in Table 1. The IEBD Rank shown for each peptide and allele was established using the netMHCIpan search engine predicting the peptides’ binding score to the respective HLA allele, whereby a lower Percentile Binding Score binding score denotes better peptide binding.

